# DeepConsensus: Gap-Aware Sequence Transformers for Sequence Correction

**DOI:** 10.1101/2021.08.31.458403

**Authors:** Gunjan Baid, Daniel E. Cook, Kishwar Shafin, Taedong Yun, Felipe Llinares-López, Quentin Berthet, Aaron M. Wenger, William J. Rowell, Maria Nattestad, Howard Yang, Alexey Kolesnikov, Armin Töpfer, Waleed Ammar, Jean-Philippe Vert, Ashish Vaswani, Cory Y. McLean, Pi-Chuan Chang, Andrew Carroll

## Abstract

Pacific BioScience (PacBio) circular consensus sequencing (CCS) generates long (10-25 kb), accurate “HiFi” reads by combining serial observations of a DNA molecule into a consensus sequence. The standard approach to consensus generation uses a hidden Markov model (pbccs). Here, we introduce DeepConsensus, which uses a unique alignment-based loss to train a gap-aware transformer-encoder (GATE) for sequence correction. Compared to pbccs, DeepConsensus reduces read errors in the same dataset by 42%. This increases the yield of PacBio HiFi reads at Q20 by 9%, at Q30 by 27%, and at Q40 by 90%. With two SMRT Cells of HG003, reads from DeepConsensus improve hifiasm assembly contiguity (NG50 4.9Mb to 17.2Mb), increase gene completeness (94% to 97%), reduce false gene duplication rate (1.1% to 0.5%), improve assembly base accuracy (Q43 to Q45), and also reduce variant calling errors by 24%.

## Introduction

Modern genome sequencing samples the genome in small, error-prone fragments called reads. At the read level, the higher error of single molecule observations is mitigated by consensus observations. In Illumina data, the consensus is spatial, through clusters of amplified molecules^1^. Pacific Biosciences (PacBio) uses repeated sequencing of a circular molecule to build consensus across time^2^. The accuracy of these approaches, and the manner they fail, ultimately limits the read lengths of these methods and the analyzable regions of the genome^3,4^

Recent breakthroughs in PacBio throughput have enabled highly accurate (99.8%) long reads (>10 kb), called HiFi reads^5^ to set new standards in variant calling accuracy^6^ and the first telomere-to-telomere human assembly^7^. The remaining sequencing errors are strongly concentrated in homopolymers^3,8^, and the need to manage these errors constrains the minimum number of passes required for acceptable accuracy, and therefore the yield and quality of PacBio sequencing.

The existing algorithm for consensus generation from HiFi sequencing data uses a hidden Markov model to create a draft consensus sequence, which is iteratively polished^9^. The underlying process of removing errors using an alignment of reads is also used in genome assembly^10^, and in assembly polishing methods like Racon^11^, Pilon^12^, and PEPPER-Margin-DeepVariant^13^. All of these methods correct from a given alignment to a reference or contig. These methods use statistical heuristics for the correction model itself, except for PEPPER-Margin-DeepVariant.

To improve consensus generation of HiFi sequencing data, we introduce a deep learning-based approach leveraging a transformer^14^ architecture. Transformers have gained rapid adoption in natural language processing^15^ and computer vision^16^. In biology, transformers have been applied to Multiple Sequence Alignment (MSA) of protein sequences^17^ and dramatically improved AlphaFold2’s protein structure prediction^18^.

We present DeepConsensus, an encoder-only transformer model that uses an MSA of the PacBio subread bases and a draft consensus from the current production method (pbccs). DeepConsensus incorporates auxiliary base calling features to predict the full sequence in a window (by default 100bp). Since insertion and deletion (INDEL) errors are the dominant class of error in this data, we train the model with a novel alignment-based loss function inspired by differentiable dynamic programming^19^. This gap-aware transformer-encoder (GATE) approach more accurately represents misalignment errors in the training process.

DeepConsensus reduces errors in PacBio HiFi reads by 41.9% compared to pbccs in human sequence data. We stratify performance across mismatches, homopolymer insertions and deletions, and non-homopolymer insertions and deletions, and DeepConsensus improves accuracy in each category. DeepConsensus increases the yield of reads at 99% accuracy by 8.7%, at 99.9% accuracy by 26.7%, and at 99.99% accuracy by 90.9%. We demonstrate that using reads from DeepConsensus improves the contiguity, completeness, and correctness of genome assembly when compared to assemblies generated using pbccs reads. Similarly, we demonstrate improved accuracy of variant calling when using DeepConsensus reads. Finally, we demonstrate that improvements in accuracy allow for longer PacBio reads lengths while retaining acceptable read accuracy, enabling improvements in contiguity of genome assembly and increasing the experimental design options for PacBio sequencing.

## Results

### Overview of DeepConsensus

An overview of the DeepConsensus algorithm is shown in Figure 1. PacBio CCS sequencing produces a set of subreads which are processed by pbccs to produce a consensus (CCS) read. Subreads are combined with the CCS read, and divided into 100 bp partitions. Each partition is then transformed into a tensor to be used as input to the DeepConsensus model for training or inference.

**Figure 1.**
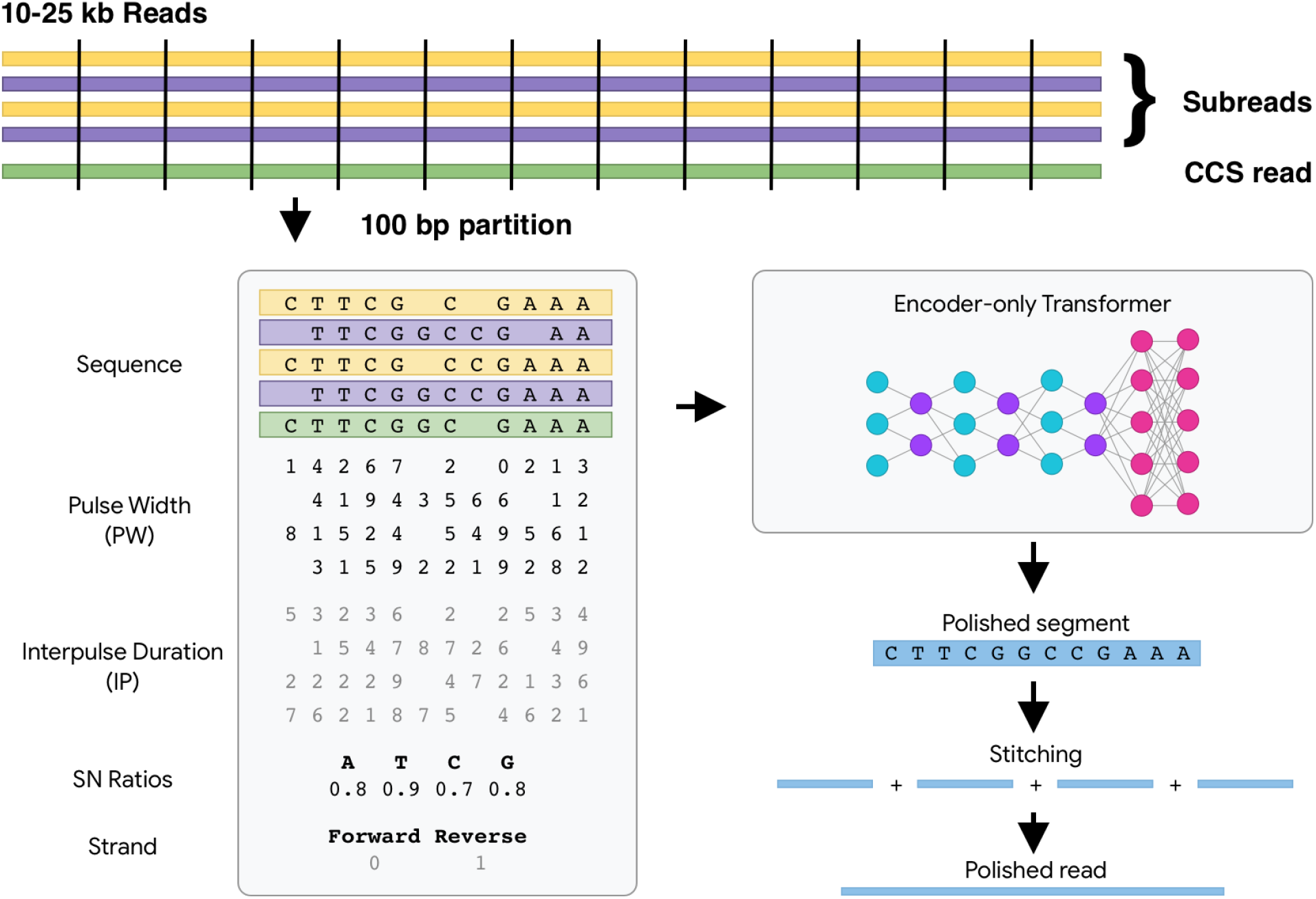
Overview of DeepConsensus. This figure illustrates the DeepConsensus workflow. Subreads are combined with a CCS read divided into 100 bp partitions. Each partition is converted to a tensor object containing the pulse width, interpulse duration, signal-to-noise (SN) ratios, and strand information. These tensors can then be used during training or inference using an encoder-only transformer. The trained model produces a polished segment which is stitched together to produce a polished read.

The tensor contains additional information beyond the sequence extracted from each subread. This includes the pulse width (PW) and interpulse duration (IP). These are raw values provided by the basecaller that are used to call bases. Additionally, DeepConsensus incorporates the signal-to-noise ratio for each nucleotide, and strand information. For training, we use a custom loss function that considers the alignment between the label and predicted sequence. For inference, the outputs for each 100bp partition in the full sequence are stitched together to produce the polished read.

### DeepConsensus increases HiFi accuracy and Yield

We first evaluated the performance of DeepConsensus (v0.1) by aligning polished 11kb chr20 reads from HG002 against a high-quality diploid assembly^20^. HiFi reads output from pbccs were processed similarly, and we used a custom script to calculate a phred-scaled read accuracy score for each read (*Q_concordance_*; See methods). When examining the intersection of reads to assess relative improvement, we observe accuracy improvements are distributed across the full range of pbccs *Q_concordance_* scores (Figure 2A). We observe an average *Q_concordance_* of 28.94 for DeepConsensus and 26.6 for pbccs, which corresponds to an average read quality improvement of *Q_concordance_* points. We also examined read accuracy by the number of subreads used to generate each HiFi read and observe *Q_concordance_* improvements for all subread bins (Figure 2B).

**Figure 2.**
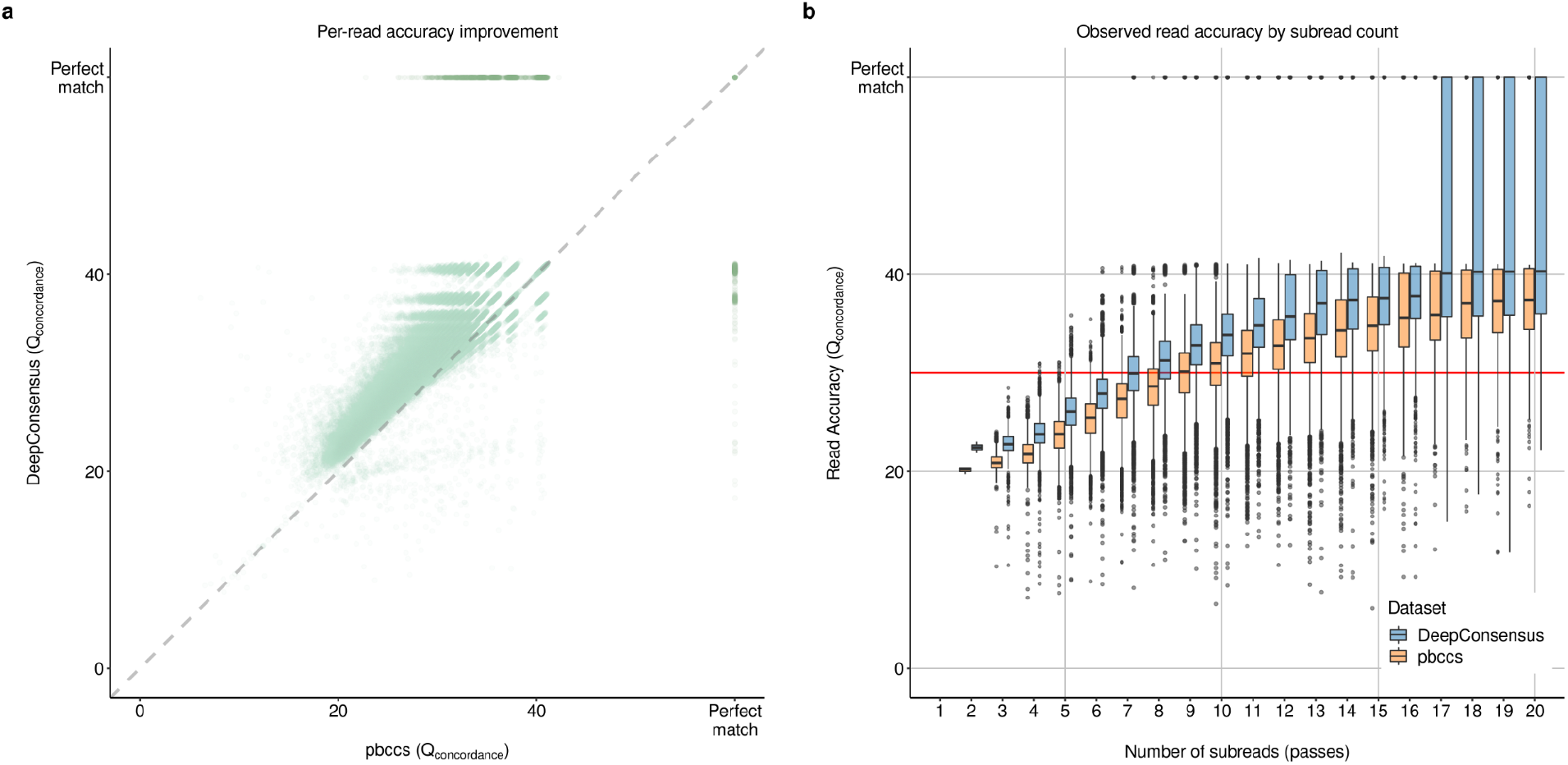
DeepConsensus improves the accuracy of CCS reads. (a) Comparison of observed read accuracy (*Q_concordance_*) for the intersection of pbccs and DeepConsensus (HG002 chr20 11kb) reads. Each light green dot corresponds to a single read. Dark green dots represent reads that were perfect matches to the reference. (b) Observed read accuracy across the number of available subreads for DeepConsensus and pbccs.

Sequencing errors can be classified by type (SNP, Indel) and according to their sequence context (homopolymer, non-homopolymer). Homopolymer indels have previously been characterized as the largest contributors to PacBio HiFi error rates^5^. We used bamConcordance^5^ to examine the improvements for each error class. Notably, DeepConsensus reduces errors across all error classes, including significant reductions in homopolymer indels and a 70.39% reduction in non-homopolymer insertions (Table 1).

**Table 1.** Corrections by error type. The average number of bp affected for each error class per-read are listed for pbccs and DeepConsensus. The percent decrease reflects the reduction in errors in DeepConsensus as compared to pbccs.

| Metric | pbccs (bp / read) | DeepConsensus (bp / read) | % Decrease |
| --- | --- | --- | --- |
| Mismatch | 1.82 | 0.99 | 45.6% |
| Homopolymer Deletion | 9.41 | 6.74 | 28.4% |
| Homopolymer Insertion | 10.28 | 5.01 | 51.3% |
| Non-Homopolymer Deletion | 1.05 | 0.97 | 7.6% |
| Non-Homopolymer Insertion | 2.06 | 0.6 | 70.9% |
| All Errors | 24.61 | 14.31 | 41.9% |

We next asked how improvements in read accuracy contribute to increases in sequencing yield. DeepConsensus and pbccs are both configured to output reads with a predicted Q > 20. We compared the total yield and yields at Q thresholds of 20, 30, 40, and perfect match. We observed that DeepConsensus increases sequencing yield across all quality bins (Table 2).

**Table 2.** Yield improvement. Polished HG002 chr20 11kb reads from pbccs and DeepConsensus are quantified according to the total number of reads, reads at given thresholds (*Q_concordance_* > 20 / 30 / 40), and reads that perfectly match the diploid assembly. Total reads is the set of initial reads output by pbccs and DeepConsensus using *Q_predicted_* > 20. The percentage increase in terms of yield achieved by DeepConsensus is listed for each category.

| Dataset | Total Reads | $Q > 20$ | $Q > 30$ | $Q > 40$ | Perfect match |
| --- | --- | --- | --- | --- | --- |
| pbccs | 90,432 | 88,561 | 47,767 | 10,207 | 4,395 |
| DeepConsensus | 103,093 | 96,260 | 60,490 | 19,485 | 9,291 |
| % Increase | 14.00% | 8.69% | 26.64% | 90.90% | 111.40% |

In addition to producing a polished sequence, our model also outputs predicted base qualities. The average base quality *Q_predicted_* should match the *Q_concordance_*. We filtered reads with identity=1 and found that the *mean*(*Q_predicted_* − *Q_concordance_*) = 2. 77. A comparison of pbccs and DeepConsensus is available in Supplementary Figure 1.

### Using DeepConsensus reads improves *de novo* assembly

To evaluate the improvements achieved in *de novo* assembly with the increased yield (Supplementary figure 2-5, Supplementary table 1) and higher quality reads from DeepConsensus, we generated phased assemblies of four human genome samples using the hifiasm^21^ assembler. We generated assemblies with reads from two SMRT Cells (HG003, HG004, HG006, HG007) and three SMRT Cells (HG003, HG004, HG006). To assess the contiguity, we derived the contig N50, NG50 and genome coverage against GRCh38 using QUAST ^22^. In Figure 3, we show the improvements in assembly quality and contiguity as a result of increased yield and quality of reads from DeepConsensus. With reads from two SMRT Cells, we see that the NG50 of the assemblies with DeepConsensus reads (17.23Mb, 12.37Mb, 31.54Mb, 8.48Mb) are on average 3x higher than assembly NG50 with pbccs reads (4.91Mb, 3.72Mb, 18.55Mb, 1.94Mb) (Figure 3a, Supplementary table 2).

**Figure 3:**
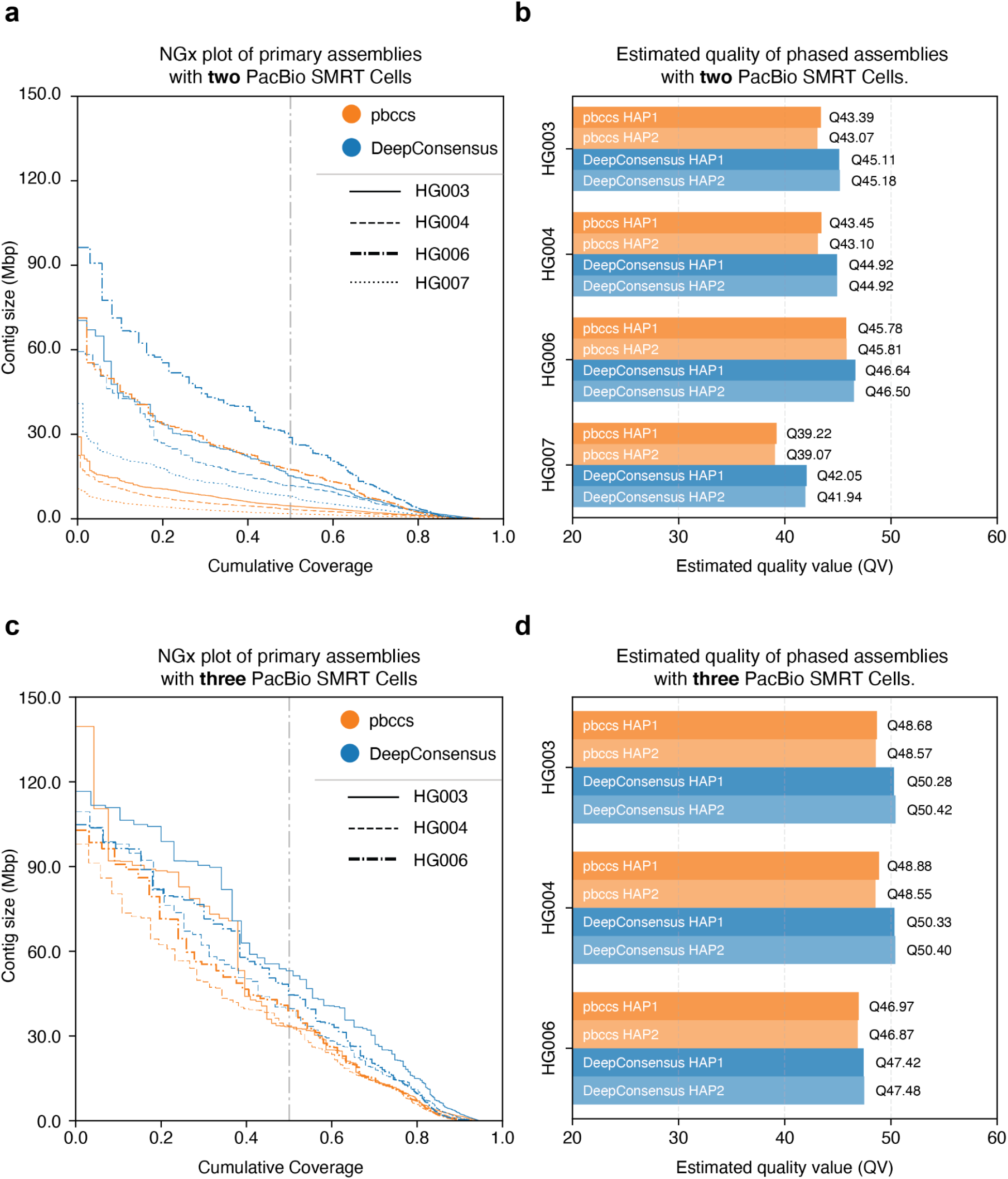
DeepConsensus improves the contiguity and quality of the genome assemblies generated with hifiasm. (a) Contiguity of the hifiasm assemblies with reads from pbccs and DeepConsensus with two PacBio SMRT Cells. (b) Reference-free estimated quality (using YAK) of the hifiasm phased assemblies with reads from pbccs and DeepConsensus with two PacBio SMRT Cells. (c) Contiguity of the hifiasm assemblies with three PacBio SMRT Cells. (d) Estimated quality of the hifiasm phased assemblies with three PacBio SMRT Cells.

We evaluated the correctness of the assembly using YAK^21^, which overlaps the assembly with *k*-mers observed in short-read sequencing. The YAK estimated quality of the assemblies with DeepConsensus reads achieve Q44 on average compared to Q42 with assemblies using pbccs reads (Figure 3b, Supplementary table 3). We also used dipcall^23^ to derive the small variants from the assembly and compared the small variants against Genome-In-a-Bottle (GIAB) truth sets^24^ of the associated sample. We observe the assemblies derived from DeepConsensus reads have on average 43% fewer total errors (false positives and false negatives) compared to the assemblies derived from pbccs reads (Supplementary table 4, 5).

To evaluate the gene completeness of the assemblies, we used asmgene^25^ with the Ensembl homo sapiens cDNA sequences as input and GRCh38 as the reference sequence. We observe that the assemblies generated with pbccs have a 2-fold higher false duplication rate (average 540 false duplications) compared to the assemblies generated with DeepConsensus (average 231 false duplications) (Supplementary table 6, 7).

Similarly, in assemblies generated with three SMRT Cells, we see that the contig NG50 of the assemblies with DeepConsensus reads (55Mb, 41Mb, 51Mb) are on average 1.3x higher than contig NG50 with pbccs reads (33Mb, 36Mb, 41Mb) (Supplementary table 2). The average assembly quality is Q49.4 with DeepConsensus reads compared to Q48.1 for assemblies with pbccs reads (Figure 3d, Supplementary table 3). The assembly-based small variant evaluation shows that assemblies from DeepConsensus reads have 33% fewer total errors compared to assemblies with pbccs reads (Supplementary table 4, 5). The gene completeness analysis shows that the assemblies generated with pbccs (average 162 false duplications) have higher number of false duplications compared to the assemblies generated with DeepConsensus (average 134 false duplications) (Supplementary table 6, 7).

In summary, we observe consistent improvements in contiguity, correctness, and completeness in assemblies generated with reads from DeepConsensus, using either two or three SMRT Cells.

### Using DeepConsensus reads improves variant calling accuracy

To assess small variant calling improvements with DeepConsensus reads, we mapped pbccs and DeepConsensus reads to the GRCh38 reference with pbmm2^25^ and called variants with DeepVariant^26^ for four human genome samples. We used the DeepVariant v1.2 PacBio HiFi model for variant calling with pbccs reads. We trained a custom DeepVariant model to call variants with DeepConsensus reads from chr1-chr19 with HG002 GIAB v4.2.1 as the small variant benchmark set.

For the variant calling analysis, we used HG003, HG004, HG006, HG007 samples. For HG003 and HG004, we used GIAB v4.2.1 and for HG006 and HG007 we used GIAB v3.3.2 benchmark set to evaluate the variants. We used hap.py^27^ to assess the variants against the GIAB benchmark set. For each sample, we report the number of false positives (FP) and false negatives (FN) variants in SNP (single nucleotide polymorphism) and INDEL (insertions and deletions) categories.

In Figure 4, we show the variant calling performance of DeepVariant with DeepConsensus and pbccs reads for two and three SMRT Cells. Variant calling with DeepConsensus reads from two SMRT Cells has on average 25% fewer errors for HG003, HG004 and 30% fewer errors for HG006, HG007 samples compared to variants with pbccs reads (Figure 4a, 4c, 4e, Supplementary Figure 6). Similarly, variants derived from DeepConsensus reads from three SMRT Cells have on average 8% fewer total errors for HG003, HG004 and 28% fewer for HG006 compared to variants with pbccs reads. Furthermore, we observe the SNP errors on average, decreasing 35% for two and 8% for three SMRT Cells of HG003 and HG004 samples (Figure 4b, 4d, 4f). Similarly, INDEL errors on average decrease 15% for two and 6% for three SMRT Cells in variants with DeepConsensus reads for HG003 and HG004 samples (Supplementary table 8). In summary, DeepConsensus improves variant calling performance across samples in both SNP and INDEL categories with reads from two and three SMRT Cells.

**Figure 4:**
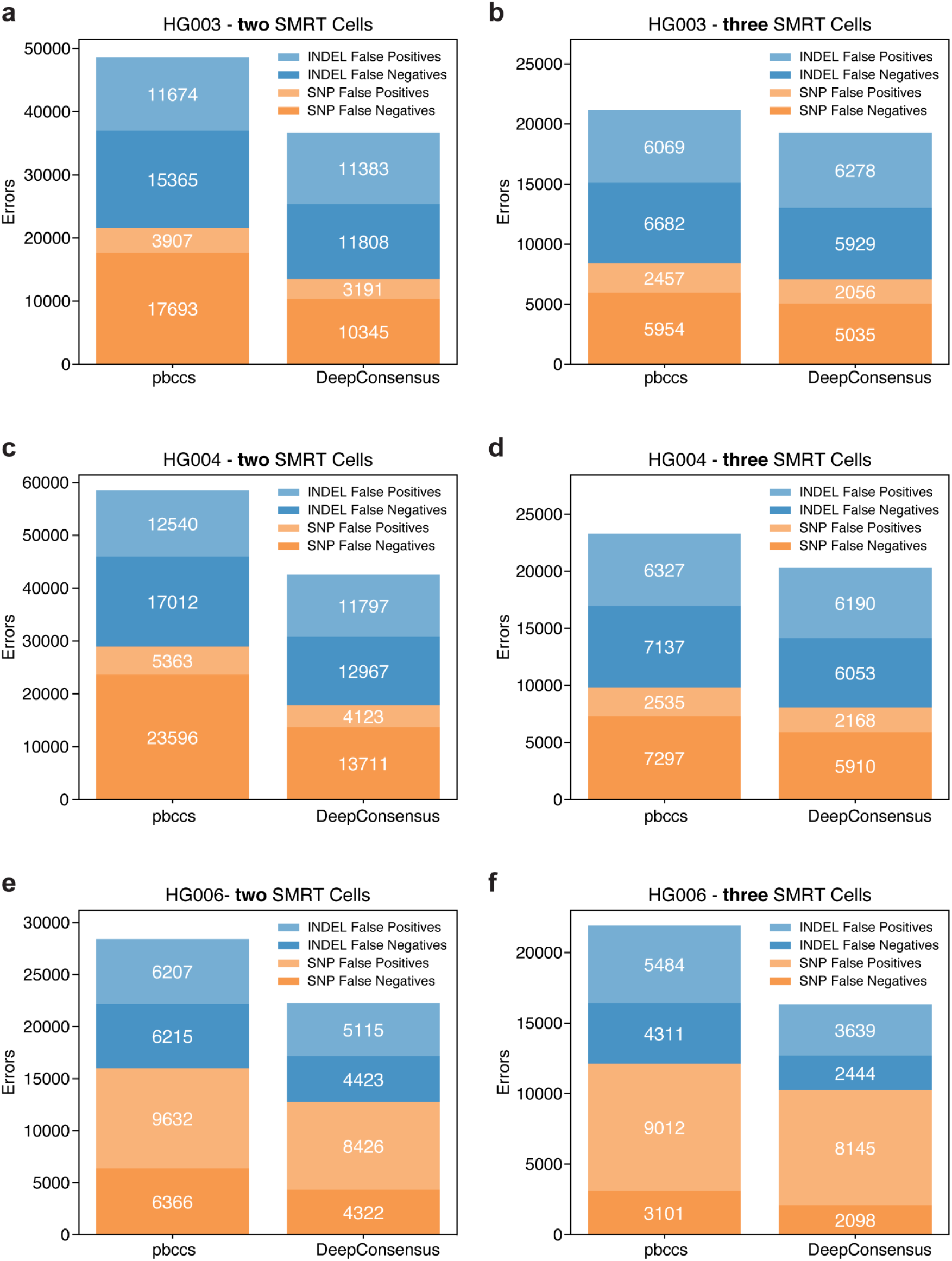
DeepConsensus improves variant calling performance of DeepVariant. (a, b) HG003 Variant calling performance of DeepVariant with pbccs and DeepConsensus reads from two and three SMRT Cells for (a, b) HG003, (c, d) HG004, (e, f) HG006.

### Use of longer reads improves yield, assembly, and variant calling

With higher consensus accuracy for HiFi reads, the number of passes can be reduced while maintaining accuracy (Figure 2b), potentially allowing for sequencing of longer insert sizes while preserving the quality of downstream analyses. To test this, we sequenced a HG002 sample with 15kb and 24kb insert sizes, each with two SMRT Cells on Sequel II System using Chemistry 2.2. We generated DeepConsensus reads for the 15kb and 24kb insert size (Figure 5a, Supplementary table 9). Details on the library preparation protocol for 24kb reads are provided in the online methods.

**Figure 5:**
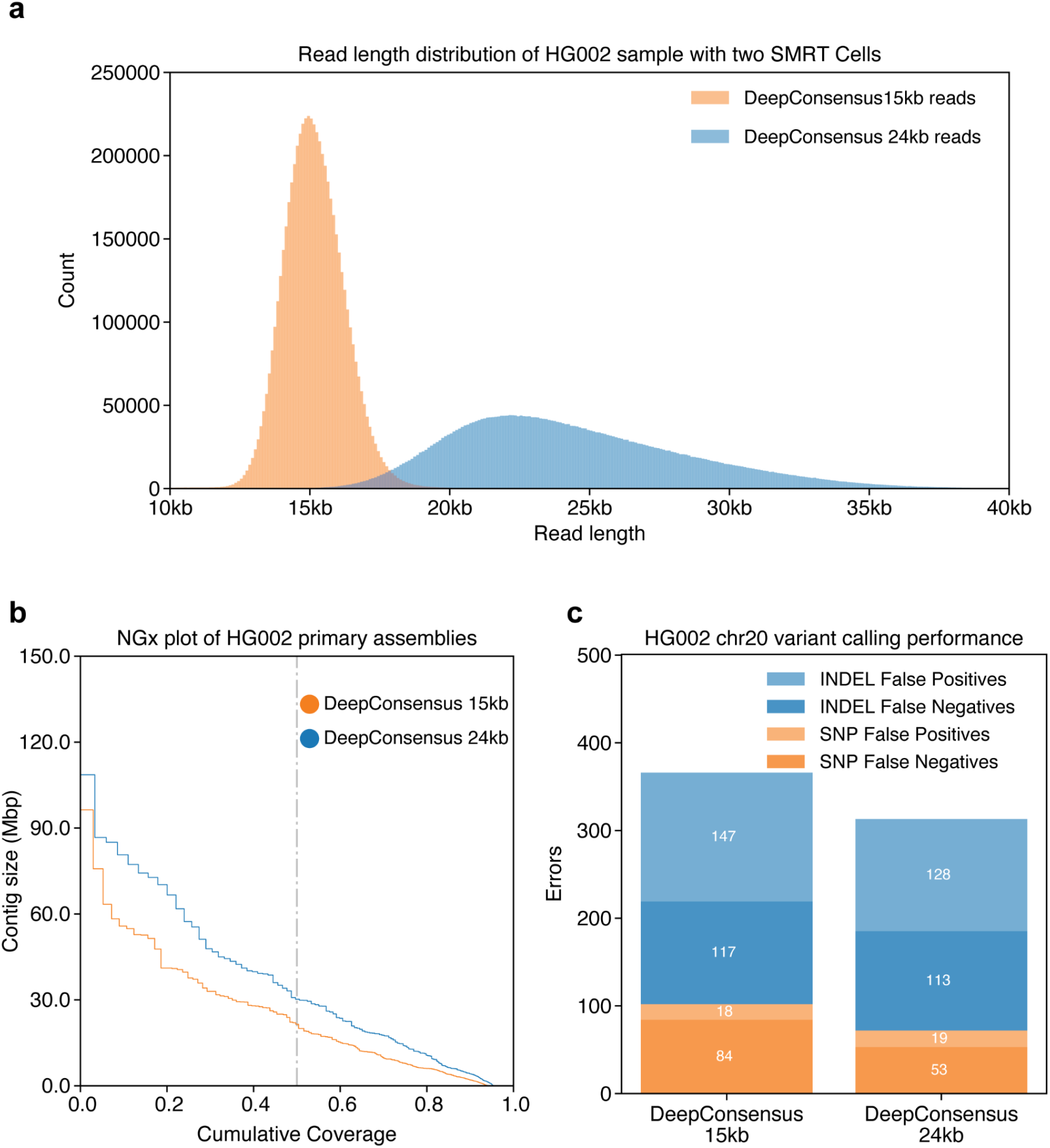
DeepConsensus with longer reads improves genome assembly contiguity. (a) HG002 read length distribution for 15kb and 24kb DeepConsensus reads from two SMRT Cells. (b) Contiguity of the HG002 hifiasm assembly with 15kb and 24kb DeepConsensus reads from two SMRT Cells. (c) HG002 variant calling performance for 15kb and 24kb reads from DeepConsensus for two SMRT Cells.

In Figure 5, we show the improvements in genome assembly and variant calling we achieve with 24kb reads compared to 15kb reads of HG002 sample. The hifiasm assembly with 24kb reads achieves higher contig NG50 of 34.05Mb compared to 24.81Mb with 15kb reads, though the assembly quality is higher with 15kb (Q51.7) reads than with 24kb (Q50.8) (Supplementary table 10, 11). The assembly-based variant calling evaluation shows the assembly with 24kb reads has higher INDEL accuracy and comparable SNP accuracy against the assembly with 15kb reads (Supplementary table 12, 13). Notably, the multi-copy gene completeness in the assembly with 24kb reads is 80.91% compared to 76.93% in the assembly with 15kb reads while the single-copy gene completeness remains comparable (97.2% with 24kb reads and 97.3% with 15kb reads) (Supplementary table 14, 15). In variant calling with DeepVariant, the 24kb DeepConsensus reads have fewer total errors compared to 15kb reads in HG002 chr20 (Figure 5c, Supplementary table 16).

In summary, the increased accuracy of DeepConsensus expands the window of experimental choices. This allows researchers to consider using longer reads for applications which disproportionately benefit, such as assembly of genomes with high duplication rates, difficult to assemble regions such as the MHC, phasing across a long gene or amplicon, or variant detection in hard-to-map regions.

## Discussion

The correction of errors in sequencing data is fundamental to both the generation of initial data from a sequencer and to downstream analyses which assemble, map, and analyze genomes^28–30^. We introduce a transformer-based consensus generation method, reducing errors in PacBio HiFi reads by 42% and increasing yield of 99.9% accurate reads by 27%. We show that with existing downstream methods, the improved reads result in better assembly contiguity, completeness, and accuracy, as well as more accurate variant calling.

The problem of correcting errors from a MSA of repeated sequencing is a single example of a broader category of problems that analyze the alignment of similar sequences. The most similar adjacent applications are error correction of Unique Molecular Identifiers^31^, as well as error correction of Oxford Nanopore Duplex reads. Genome assembly polishing, which uses alignments of sequences from many molecules, is a similar application^11,13,32^. DeepConsensus models could be trained for these applications with minimal changes to its architecture. The gap-aware loss function used in the GATE approach could have utility to broader MSA-related problems. For example, related work by Rao 2021^17^ demonstrated improved prediction performance across multiple tasks, including contact maps and secondary structure, and Avsec et al. 2021^33^ used a long-range Enformer to predict gene expression. These applications could potentially benefit from the incorporation of alignment-based loss used in DeepConsensus, or the DeepConsensus framework could be applied to similar problem areas.

DeepConsensus presents opportunities to alter experimental design to better leverage its improvements to accuracy. We demonstrate that DeepConsensus allows for longer read lengths while maintaining a high standard of read accuracy and yield. Certain applications, such as assembling difficult genome regions, may disproportionately benefit from use of longer reads. Additionally, because DeepConsensus learns its error model directly from training data, it allows a tighter coupling between library preparation, instrument iteration and informatics. DeepConsensus could be trained on data from a modified procedure or additional data stream to more accurately estimate the potential advantage of the new method, decreasing the chance that the modification’s advantages might not be apparent due to optimization of the informatics to the older approach.

The improvements we demonstrate to assembly and variant calling use unmodified downstream tools (hifiasm), or tools with unmodified heuristics that use an adapted model (DeepVariant). Further iterating on the heuristics in these methods may allow them to take additional advantage of the DeepConsensus error profile, or better use its higher yield of longer reads.

Future improvements to DeepConsensus include training with an expanded dataset that includes additional samples and chemistries, since our current training datasets only include Sequel II data from a few SMRT Cells. Assessing DeepConsensus on non-human species and, if needed, supplementing training data with diverse species is an area of active development. There are substantial opportunities for improvements by refining the attention strategy, for example AlphaFold2 uses a modified axial attention^34^, or by leveraging efficiency improvements to the transformer self-attention layer to consider wider sequence contexts^35–37^. Investigating the trade-offs between model size and accuracy could also enable faster versions which preserve high accuracy. These and other improvements will enable DeepConsensus to help scientists realize the potential yield and quality of their sequencing instruments and projects.

## Code Availability

Code and pretrained models are available on https://github.com/google/deepconsensus.

## Data Availability

Sequencing data, predictions, and analysis files are available at: https://console.cloud.google.com/storage/browser/brain-genomics-public/research/deepconsensus/publication

Sequencing data is available from the following sources:

- Sequel II data from Novogene: https://console.cloud.google.com/storage/browser/brain-genomics-public/research/sequencing
- 15kb HG002 and 24kb HG002 reads from PacBio: https://console.cloud.google.com/storage/browser/brain-genomics-public/research/deepconsensus/publication/sequencing
- Sequel II data from PacBio: https://downloads.pacbcloud.com/public/dataset/HG002_SV_and_SNV_CCS/
- HG002 diploid assembly: https://obj.umiacs.umd.edu/marbl_publications/hicanu/hg002_hifi_hicanu_combined.fasta.gz

## Online Methods

### Generation of 24kb PacBio reads

DNA was extracted from HG002/NA24385 cell pellets (Coriell Institute) with the MasterPure Complete DNA and RNA Purification Kit (Lucigen MC85200) and sheared on Megaruptor3 (Diagenode) at speed 30. SMRTbell libraries were constructed with SMRTbell Express Template Prep Kit 2.0 (PacBio 100-038-900). Size selection was performed with BluePippin (Sage Science) with an 18kb high-pass filter. Sequencing was performed on the Sequel II System using Chemistry 2.2 and 30 hour movies.

### Dataset preparation

For all SMRT Cells, we ran pbccs on the subreads to generate CCS sequences. pbccs generates a prediction for the overall read quality for each CCS read, and reads below Q20 are filtered out of the final HiFi read set. For dataset generation, we did not apply any filtering based on read quality for the CCS reads, and reads of qualities were included for training and inference. To generate labels for each set of subreads, the CCS sequence predicted by pbccs was mapped to the HG002 diploid assembly. The coordinates of the primary alignment were used to extract the label sequence from the HG002 diploid assembly.

Subreads and labels were aligned to the corresponding CCS sequence. The cigar string from this alignment was used to match bases across the subreads and assign a label for each position. Subreads were broken up into 100bp windows, and the corresponding label for each window was extracted from the full label sequence. In some cases, the label was longer than the subreads due to bases for which there was no support in the subreads.

Each subread base has associated pulse width (PW) and interpulse duration (IPD) values, and each set of subreads has four signal to noise ratio (SN) values, one for each of the four canonical bases. PW and IP values were capped at 9, and SN values were rounded to the closest integer and capped at 15.

### Model and training

The Transformer has emerged as the primary architecture for language understanding and generation tasks^14,15^. It uses self-attention to efficiently capture long and short-range interactions between words, crucial for understanding text. In recent work, this capability has been successfully leveraged to improve modeling of protein sequences^38^.

We train a six layer encoder-only transformer model with a hidden dimension of 560 and 2 attention heads in each self-attention layer. The inner dimension of the feedforward network in each encoder layer is 2048. The model considers 100 bases at a time from the full subreads, and the input at each position contains subread sequences and auxiliary features. The maximum number of subreads considered is 20. Auxiliary features include the pulse width (PW) and interpulse duration (IPD) measured by the basecaller, the signal to noise ratio (SN) for the sequencing reaction, the strand of each subread, and the sequence of the CCS read as predicted by pbccs. Each feature type is embedded using a separate set of learned embeddings, which are trained jointly with the model. An embedding size of two is used for the subread strand, and all other embeddings are of size eight. We used positional encodings that were a mix of sampled sine and cosines, as defined in the Transformer^14^. For training, the Adam optimizer was used with a learning rate of 1e-4, and input, attention, and ReLU dropout values were set to 0.1. Our implementation builds off the one provided in the [Tensorflow Model Garden](https://github.com/tensorflow/models/blob/master/official/nlp/transformer/transformer.py).

For some examples, there exists a base in the label for which there is no evidence in any of the subreads. The predicted sequence for such examples would be longer than the input sequence length. The transformer encoder block outputs an encoding for each input token. In natural language applications, variable-length prediction is implemented using a decoder block, which is not constrained in the number of outputs. For consensus generation, we did not use a decoder block due to computational constraints. To allow for variable-length prediction using only the encoder, we add a fixed number of padding tokens to the input sequence for each window. This allows the model to predict sequences longer than the subread sequences by replacing some of the padding tokens with additional bases.

The outputs from the encoder block are independently decoded using a shared feedforward layer with softmax activation. At each position, we predict a distribution over the vocabulary, which consists of the four canonical bases, A, C, T, G, and an additional token to represent alignment gaps or padding, which we denote as $.

For training, we used chromosomes 1-19 from PacBio Sequel II sequencing of HG002, an extensively characterized genome curated by Genome in a Bottle^39^. Chromosomes 21 and 22 were used for tuning model parameters, and chromosomes 19 and 20 were held out entirely during training and used for final assessment. For additional full holdouts, we use PacBio Sequel II sequencing of HG003, HG004, HG006, HG007. Models were trained for 50 epochs on 128 TPU v3 cores TPUs with a batch size of 256 for each core. Five models were trained with the production settings, and we chose the checkpoint with lowest loss on the tuning data. A custom gap-aware alignment loss was used, which is described in more detail in the following section. We call the combination of the gap-aware loss with transformer-encoder architecture GATE (gap-aware transformer-encoder).

### Loss Function

Given an input MSA consisting of subreads and a consensus read and auxiliary features, the output of the transformer is a sequence *y* = *y*_1_ *y*_2_ … *y_N_* of probability distributions over the 5-letter alphabet *N* = {*A*, *T*, *C*, *G*, $}, where $ refers to an empty character to model possible insertion errors in the HiFi consensus or padding. In other words, each *y_i_* is a probability distribution of nonnegative entries that satisfies *y_i_* (*A*) + *y_i_* (*T*) + *y_i_* (*C*) + *y_i_* (*G*) + *y_i_* ($) = 1. At inference time, the predicted nucleotide sequence *z* = *z*_1_ *z*_2_ … *z_N_* is simply obtained by keeping the character with largest probability at each position, i.e., *z_i_* = *argmax*_*a*∈*N*_ *y_i_* (*a*) and removing the $ character from the resulting sequence. At train time, when parameters of the transformer-based model are updated, we need to define a loss function *loss*(*y*, *t*) differentiable with respect to the transformer output *y* given the correct nucleotide sequence *t* = *t*_1_ *t*_2_ … *t_M_* (notice that the lengths *N* of the transformer output and *M* of the correct nucleotide sequence may differ due to possible insertion or deletion in the consensus read). If we know that a given position 1 ≤ *i* ≤ *N* of the transformer output should predict the nucleotide at position 1 ≤ *j* ≤ *M* of the true sequence, then it is natural to use the cross-entropy loss *loss_CE_* (*y_i_*, *t*) =− log *y_i_* (*t_j_*) to assess how good the prediction is. However, we need to choose which position of *y* predicts which position of *t*. For that purpose, we formally define an *alignment* of length *k* as an increasing subset of *k* positions π = {1 ≤ π(*y*, 1) < π(*y*, 2) <… < π(*y*, *k*) ≤ *N*, 1 ≤ π(*t*, 1) < π(*t*, 2) <… < π(*t*, *k*) ≤ *M*} in both *y* and *t*, such that position π(*y*, *ν*) in *y* predicts position π(*t*, *ν*) in *t*, for *ν* = 1,…, *k*. Given such an alignment, positions 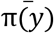 of *y* not in the alignment correspond to *insertion* errors, and ideally the prediction in those positions should be $ so that they are removed from the prediction at test time. For those positions, we therefore use the cross-entropy loss *loss_CE_* (*y_i_*, $). On the other hand, positions 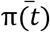 of *t* not in the alignment corresponds to *deletion* errors, i.e., nucleotides in the correct sequence that are missed in the MSA. For those errors, we consider a fixed error γ > 0, which is a parameter to be tuned. In total, given an alignment π, the total loss is defined as the sum of cross-entropy losses over aligned positions and insertion/mutation losses:

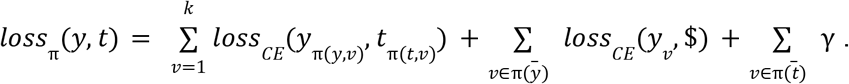

This loss depends on the arbitrary alignment π, which ideally should be chosen as a function of *y* and *t* so that the total loss is small. We therefore finally define the alignment loss as a (smooth) minimum over 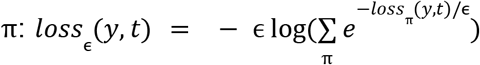, where ϵ ≥ 0 is a parameter to control how suboptimal alignments contribute to the loss. At the limit ϵ = 0, we simply keep the best alignment 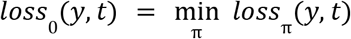, and taking ϵ > 0 allows us to create a smoother loss function to better align *y* and *t*. This loss is a particular case of the losses studied previously^19^, and we follow their approach to derive an efficient implementation to compute the loss and its gradient in *y* using differentiable dynamic programming, with a specific wavefront formulation to accelerate the computation on GPUs and TPUs.

### Output FASTQ generation

DeepConsensus predictions for each 100bp window are joined together and $ tokens are removed to produce the final sequence that is output to FASTQ. Predicted base quality scores are generated from the output distribution at each position. The raw quality score for each base, *q_i_*, is computed as follows, where *y_i_* is the output distribution at position *i*: *q_i_* = − 10 *log*_10_ (1 − *max*(*y_i_*)). Each raw quality score is rounded to the closest integer and capped at a maximum value of 60 to produce the final base quality score, *Q_i_*. Final base qualities are used to compute an overall read quality, *Q_pred_*, in the following calculation, which sums over all positions in the predicted sequence: 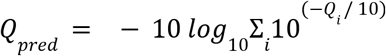. Reads with an overall predicted quality above 20 were written to the final output FASTQ, along with the corresponding quality string.

### Analysis Methods

#### Assessing read accuracy

HG002 11kb predictions were mapped to a high-quality HG002 diploid assembly^20^. For each primary alignment, the calculate_identity_metrics.py [link] script was used to compute identity which is defined below.

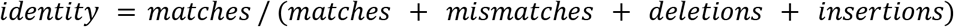

The read identity values are used to compute the concordance read qualities, *Q_concordance_*, which are computed as phred-scaled scores of the identity: *Q_concordance_* = − 10 *log*_10_(1 − *identity*).

Reads with identity scores of 1 are separately categorized as having a ‘perfect match.’ Subread counts were determined using the np tag (number of full-length passes). The np tag was extracted from the consensus reads BAM output by pbccs.

We also use the [bamConcordance tool](https://github.com/PacificBiosciences/hg002-ccs/blob/master/concordance/bamConcordance), which reports the concordance between a read and a reference sequence along with error counts for each read. Error counts are broken down into five categories: mismatches, homopolymer insertions and deletions, and non-homopolymer insertions and deletions. We use the bamConcordance output to assess the quality of reads and calculate the percentage error reduction across different categories.

#### Generating phased diploid assemblies with hifiasm

We used hifiasm version 0.15.3-r339 to generate phased assemblies. We used the default hifiasm parameters which have duplication purging on for the phased assemblies. We converted the primary assembly graph to get the primary assembly sequence and each of the haplotype graphs to generate the assembly sequences for each haplotype. Detailed execution parameters and commands are provided in the supplementary notes.

#### Reference free assembly quality estimation with YAK

We used YAK version 0.1-r62-dirty to derive estimated quality of the assemblies. For each sample, we generated a k-mer set with *k*=31 from Illumina short-reads of the same sample. Then we ran YAK to determine the quality of each haplotype sequence we generated during the hifiasm assembly generation process. YAK reports a Q value for assemblies which is a Phred-scale contig base error-rate derived by comparing 31-mers in contigs and 31-mers in the short reads of the same sample. We report the *balanced_qv* value reported by YAK as the estimated quality value of the assembly. The parameters and commands used are provided in the supplementary notes.

#### Assembly-based small variant calling assessment using dipcall

We used dipcall version 0.3 to derive small variants from the phased assemblies. Dipcall aligns the contigs to a reference sequence and derives a set of variants from the contig to reference alignment. We then compared the derived small variants against the Genome-In-a-Bottle truth set of the associated sample. For all male samples we used `-x hs38.PAR.bed` parameter as suggested in the documentation of dipcall.

To assess the small variants derived from the HG002, HG003 and HG004 sample we used GRCh38 as reference and GIAB_v4.2.1 as the truth set for small variants. For HG005, HG006, HG007 samples, we used GRCh37 and GIAB_v3.3.2 as the truth sets. All truth sets are the latest available truth set from GIAB for the associated sample. We used hap.py to assess the quality of the variant calls. Commands and parameters used to run dipcall are provided in the supplementary notes.

#### Gene completeness assessment with asmgene

We used asmgene version v2.21 to determine the gene completeness of the assemblies. First, we aligned the [Ensembl cDNA sequences](http://ftp.ensembl.org/pub/release-102/fasta/homo_sapiens/cdna/Homo_sapiens.GRCh38.cdna.all.fa.gz) release 102 to the GRCh38 reference genome using minimap2 (v2.21) and found 35374 single-copy genes and 1253 multi-copy genes in the reference. Then for each sample, we aligned the sample cDNA sequences to each of the haplotype sequence of the assemblies and derived how many single-copy genes remained single copy (`full_sgl` reported by asmgene) and how many were duplicated (`full_dup` reported by asmgene). Similarly, we reported how many multi-copy genes remained multi-copy in the assembly (`dup_cnt` reported by asmgene). We derived the following metrics to assess the gene completeness of the assemblies:

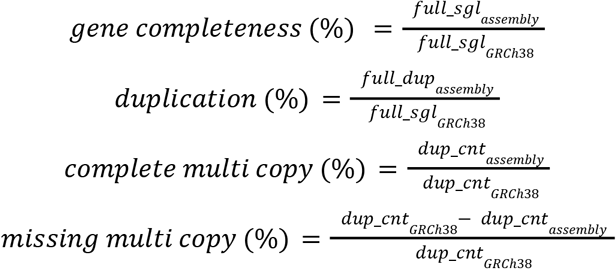

Detailed commands and parameters of asmgene are provided in the supplementary materials.

#### Assembly statistics with QUAST

We used QUAST version v5.0.2 to derive assembly N50, NG50, total assembly size and genome completeness of the assembly. QUAST is a reference-based assembly evaluation method that uses a reference sequence of the same sample or to determine the quality of the assembly. For our analysis, we used GRCh38 as the reference for each assembly.

We used N50 which is the sequence length of the shortest contig at 50% of the total assembly length to determine contiguity of the assembly. NG50 is the sequence length of the shortest contig at 50% of the estimated genome length. For our human genome assemblies, we used GRCh38 as the reference sequence so we used 3272116950bp (3.2gb) as the estimated genome length to derive NG50. We only report N50, NG50, total assembly length and genome completeness against GRCh38 from the QUAST report. Detailed parameters and commands are provided in the supplementary notes.

#### Variant Calling

DeepVariant performs variant calling in three stages: make_examples, call_variants, and postprocess_variants. The make_examples stage identifies candidate variants and generates input matrices containing pileup information. call_variants runs the input matrices through a neural network model, and the postprocess_variants converts the neural network outputs to a variant call and outputs a VCF.

We used the latest DeepVariant model for PacBio HiFi data, v1.2, to call variants in pbccs predictions. Polished DeepConsensus reads or pbccs HiFi reads were aligned to GRCh38. This model was fine tuned from the Illumina WGS v1.2 model using PacBio HiFi sequencing reads generated using pbccs. Since DeepConsensus reads display different error characteristics than pbccs reads, we fine tuned a new DeepVariant model for DeepConsensus. This model was also initialized from the v1.2 Illumina WGS model, and the training data consisted of 11kb and 24kb Sequel II reads for HG002. We mixed both phased and unphased reads for the training, similar to what is done for the v1.2 PacBio model. Chromosomes 1-19 were used for training, chromosomes 21-22 were used for tuning, and chromosome 20 was held out entirely.

## Author contributions

GB, PCC, and AC conceived the study. GB, DEC wrote DeepConsensus and trained models. GB, DEC, KS, TY, MN performed experiments with DeepConsensus reads and made figures and documentation. FLL, QB, JPV conceived and implemented the alignment loss strategy, which DEC integrated into DeepConsensus. AMW, WJR, AT provided insight into PacBio data, identified areas for improvement, suggested informative features, and provided code for pre-processing and evaluation. WA experimented with embedding strategies. AK and AT contributed to efficient processing of PacBio reads. HY coordinated data acquisition and research agreements. JPV, AV, CYM, PCC, and AC provided guidance on experimental design, architecture, and code review. GB, DEC, KS, TY, FLL, QB, AMW, WJR, MN, JPV, AV, CYM, PCC, AC wrote the paper.

