## Supplementary Material for "DeepConsensus: Gap-Aware Sequence Transformers for Sequence Correction"

### Supplementary Notes

#### Software commands

##### Generating DeepConsensus input files with pbmm2 and samtools

We started with the subreads BAM and CCS reads BAM files output by pbccs. Samtools v1.1 was used to convert the CCS reads BAM to a FASTA:

```
samtools fasta "ccs.bam" > "ccs.fasta"
```

We used pbmm2 v1.4.0 for mapping subreads to CCS reads using the following command:

```
pbmm2 align --sample "sample" --preset SUBREAD --sort "ccs.fasta"
"subreads.bam" "aligned.subreads.bam"
```

Only subread alignments to the correct molecule were retained. We used samtools and awk to filter incorrect alignments using the below command:

```
samtools view -h "aligned.subreads.bam" | \
  awk '{ if($1 ~ /^@/) { print; } else { split($1,A,"/"); \
  split($3,B,"/"); if(A[2]==B[2]) { split(A[3],C,"_"); \
  print $0 "\tqs:i:" C[1]; } } }'
```

##### Running inference with DeepConsensus

DeepConsensus v0.1 was run with the following command:

```
python3 -m deepconsensus.scripts.run_deepconsensus \
  --input_subreads_aligned=subreads.aligned.bam \
  --input_subreads_unaligned=subreads.bam \
  --input_ccs_fasta=ccs.fasta \
  --output_directory=deepconsensus_output \
  --checkpoint=model-50
```

The outputs are sharded FASTQs, which are concatenated together with cat to produce the final FASTQ.

##### Aligning DeepConsensus predictions with pbmm2

DeepConsensus predictions and CCS reads from pbccs were mapped to the HG002 diploid assembly or reference genome using the below command. The `--unmapped` flag was added to the below command for assembly and variant calling analysis pipelines:

```
pbmm2 align --sample "sample" --preset HIFI --sort -c 0 -y 70
"ref.fa" "query.reads.fasta" "aligned.reads.bam"
```

#### Generating phased diploid assemblies with hifiasm

HiFiasm (version 0.15.3-r339) was run with the following command to generate the diploid human genome assemblies:

```
hifiasm -o "outputPrefix" -t "nThreads" "reads.fastq"
```

We ran the following commands to convert the output GFA files to fasta files.
For the primary assembly we used:

```
awk '/^S/{print ">"$2;print $3}' "outputPrefix.bp.p_ctg.gfa" >
"outputPrefix.bp.p_ctg.fa"
```

For the haplotype-resolved GFA files for the diploid sample, we used the following commands to get haplotype-resolved assembly:

```
awk '/^S/{print ">"$2;print $3}' "outputPrefix.bp.hap1.p_ctg.gfa" >
"outputPrefix.bp.hap1.p_ctg.fa"

awk '/^S/{print ">"$2;print $3}' "outputPrefix.bp.hap2.p_ctg.gfa" >
"outputPrefix.bp.hap2.p_ctg.fa"
```

#### Reference free assembly quality estimation with YAK

We used yak (version 0.1-r62-dirty) to derive estimated consensus accuracy (Q) of the assemblies.

To evaluate the assembly of a sample we first built the k-mer database from Illumina paired-end short reads of the same sample using this command:

```
yak count -b37 -t32 -o "KMER_DB" <(zcat sr*.fq.gz) <(zcat sr*.fq.gz)
```

Then we estimated the quality of the phased assembly using the following command:

```
yak qv -t "nThreads" -p -K 3.2g -l 100k "KMER_DB"
"outputPrefix.bp.hap1.p_ctg.fa" > "outputPrefix.hap1.yak.qv.txt"

yak qv -t "nThreads" -p -K 3.2g -l 100k "KMER_DB"
"outputPrefix.bp.hap2.p_ctg.fa" > "outputPrefix.hap2.yak.qv.txt"
```

From the outputs `"outputPrefix.hap1.yak.qv.txt"` and `"outputPrefix.hap2.yak.qv.txt"` we reported the balanced error as the quality of the assembly we evaluated.

#### Assembly-based small variant calling assessment using dipcall

We used dipcall (version 0.3) to derive small variants from the phased assemblies.

For male samples such as HG002, HG003, HG006, we used the following command:

```
dipcall.kit/run-dipcall -x dipcall.kit/hs38.PAR.bed
"outputPrefix.dipcall_output" "reference"
```

```
"outputPrefix.bp.hap1.p_ctg.fa" "outputPrefix.bp.hap2.p_ctg.fa" >
dipcall_instructions.mak

make -j2 -f dipcall_instructions.mak
```

For female samples such as HG004, HG007 we used the following command:

```
dipcall.kit/run-dipcall "outputPrefix.dipcall_output" "reference"
"outputPrefix.bp.hap1.p_ctg.fa" "outputPrefix.bp.hap2.p_ctg.fa" >
dipcall_instructions.mak

make -j2 -f dipcall_instructions.mak
```

From the output we compared `outputPrefix.dipcall_output.dip.vcf.gz` variant call set against GIAB truth set using hap.py small variant evaluation program. For HG002, HG003 and HG004 we used GIAB v4.2.1 benchmarking set and GRCh38 as the reference and for HG006, HG007 we used GIAB v3.3.2 benchmarking set and GRCh37 as the reference. The commands for hap.py is provided in the associated section.

#### Gene completeness assessment with asmgene

We assessed the gene completeness of the phased assemblies with asmgene (version v2.21). We first aligned the [Ensembl cDNA sequences] to the GRCh38 reference genome using minimap2 (v2.21) using the following command:

```
minimap2 -cxsplice:hq -t32 "GRCh38.fa" "homo_sapiens_cdna.fa" >
"asmgene_output_prefix.ref.cdna.paf"
```

Then we align the cDNA sequences to each haplotype of the phased assemblies.

```
minimap2 -cxsplice:hq -t32 "outputPrefix.bp.hap1.p_ctg.fa"
"homo_sapiens_cdna.fa" > "asmgene_output_prefix.HAP1.cdna.paf"

minimap2 -cxsplice:hq -t32 "outputPrefix.bp.hap2.p_ctg.fa"
"homo_sapiens_cdna.fa" > "asmgene_output_prefix.HAP2.cdna.paf"
```

Then we use asmgene tool available from paftools.js to derive the gene completeness metrics for each haplotype.

```
paftools.js asmgene -a "asmgene_output_prefix.ref.cdna.paf"
"asmgene_output_prefix.HAP1.cdna.paf" >
"asmgene_output_prefix.hap1.genestats"

paftools.js asmgene -a "asmgene_output_prefix.ref.cdna.paf"
"asmgene_output_prefix.HAP2.cdna.paf" >
"asmgene_output_prefix.hap2.genestats"
```

#### Assembly statistics with QUAST

We used QUAST (version v5.0.2) to derive the assembly size, N50 and NG50 metrics of the assembly against the GRCh38 reference sequence. For the assembly size, N50 and NG50 values, we used the primary assembly we get from hifiasm assembler. We run the following command to derive the assembly metrics:

```
python /root/tools/quast/quast-5.0.2/quast-lg.py -t "nThreads" -o
"QUAST_output_directory" -r "GRCh38.fa" --large
outputPrefix.bp.p_ctg.fa
```

Within the output directory the `report.txt` file contains the assembly size, N50 and NG50 numbers that we report in our evaluation.

#### Variant calling with DeepVariant

We ran DeepVariant with the provided convenience script to call variants. Here is the command used:

```
/opt/deepvariant/bin/run_deepvariant --model_type="PACBIO"
--use_hp_information --ref="ref.fa" --reads="reads.bam"
--output_vcf="output.vcf" --num_shards="$(nproc)"
```

For the pbccs baseline, we used the latest DeepVariant model for PacBio data, v1.2. For DeepConsensus, we trained a custom model which was specified using the `--customized_model` in the above command.

Variant calls were evaluated using v0.3.12 of hap.py from the jmcdani20/hap.py Docker image. The command used was:

```
/opt/hap.py/bin/hap.py "truth.vcf" "output.vcf" -f "truth.bed" -r
"ref.fa" -o "output/happy.output"
--stratification="v2.0-GRCh38-stratifications.tsv" --engine=vcfeval
--pass-only
```

Truth VCF and BED files come from the Genome in a Bottle (GIAB) truth sets. The v4.2.1 truth set was used for HG002, HG003, and HG004, and the v3.3.2 truth set was used for HG006 and HG007.

### Supplementary results

#### Supplementary figures

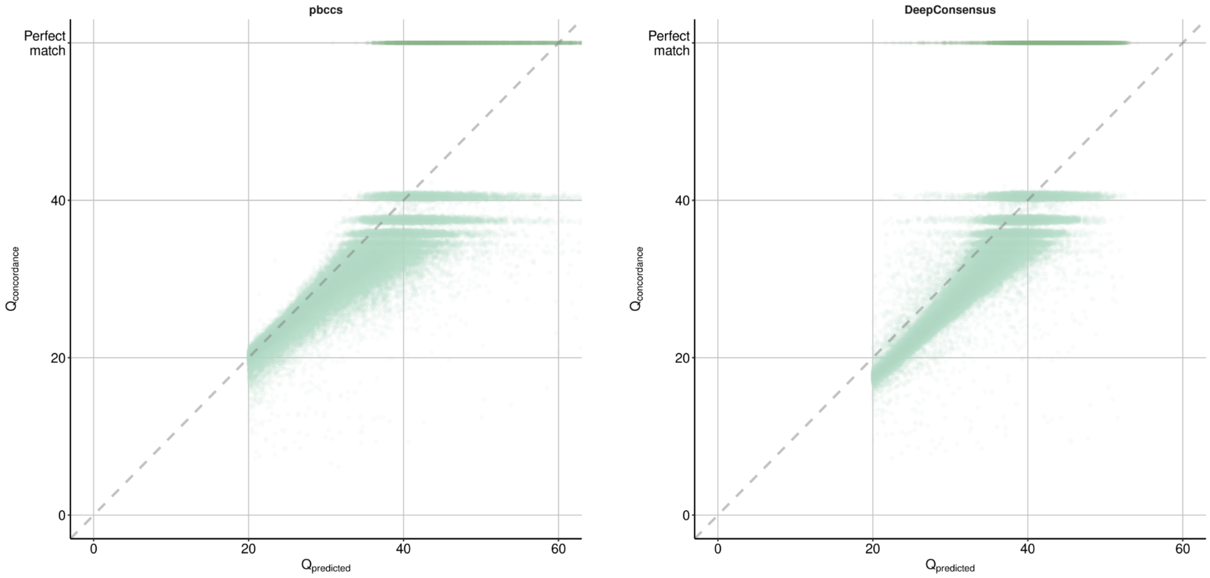


**Supplementary Figure 1** The observed read quality ($Q_{concordance}$) and predicted read quality ($Q_{predicted}$) are plotted against one another for both pbccs and DeepConsensus. Reads are from HG002 chr20 and have a length of 11kb.

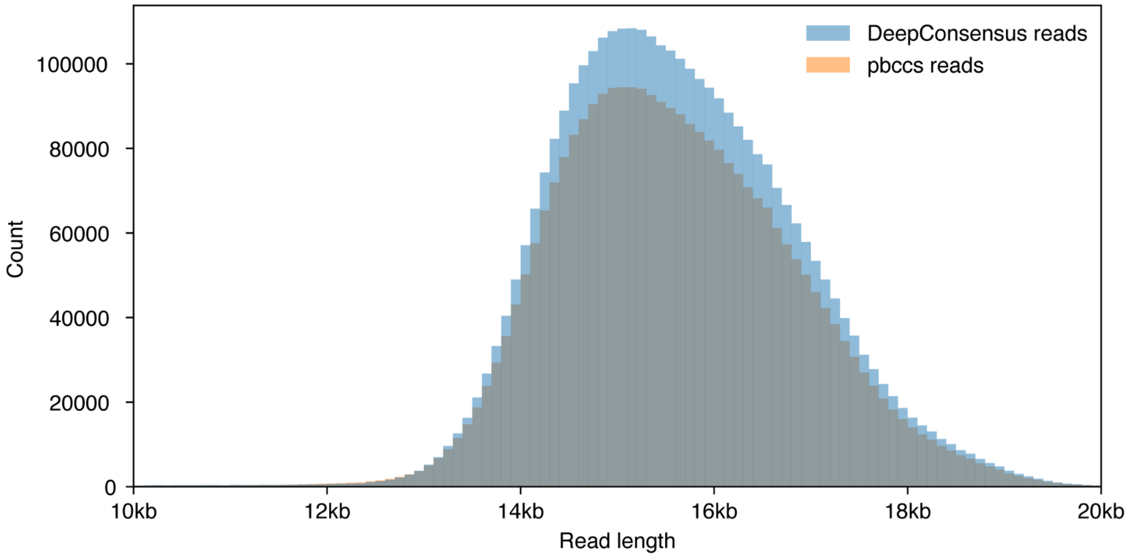


**Supplementary Figure 2** Read length distribution of HG003 two SMRT Cells reads from pbccs and DeepConsensus.

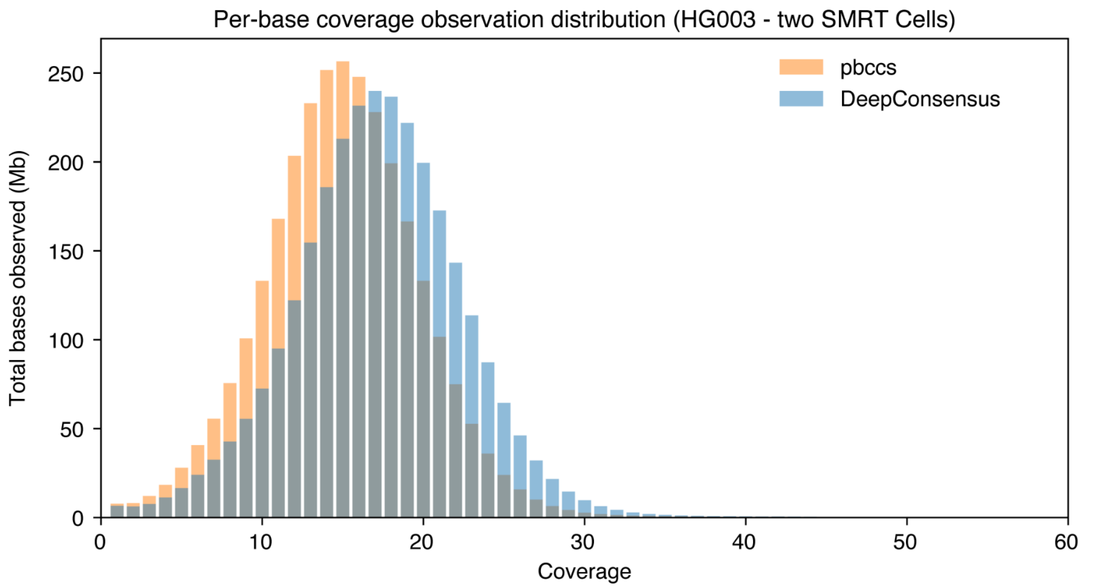


**Supplementary Figure 3** Distribution of observed coverage of HG003 two SMRT Cells reads from pbccs and DeepConsensus.

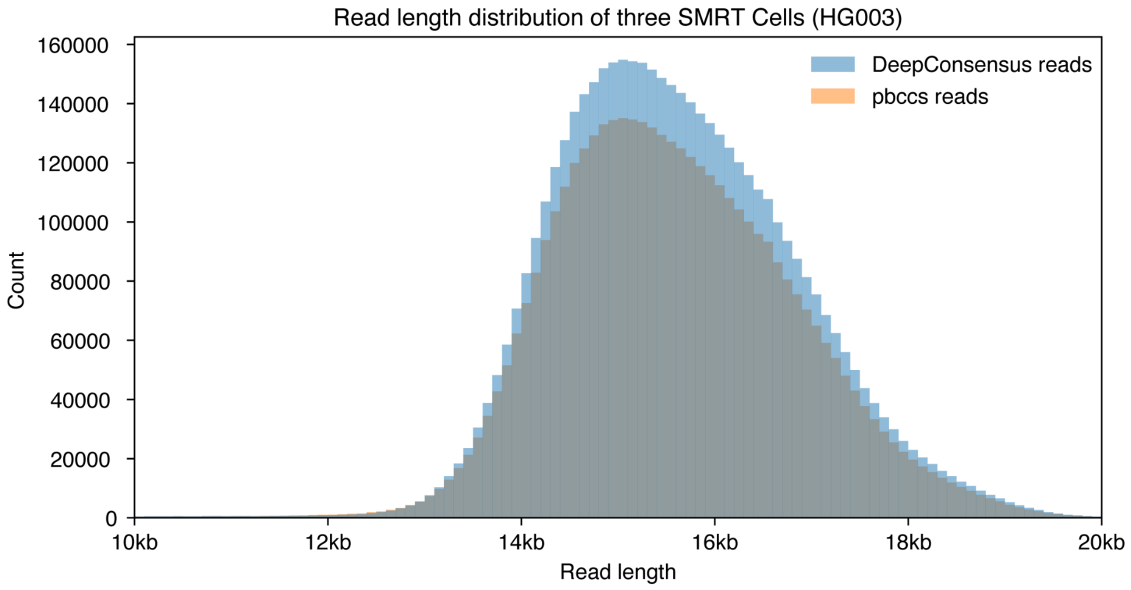


**Supplementary Figure 4** Read length distribution of HG003 three SMRT Cells reads from pbccs and DeepConsensus.

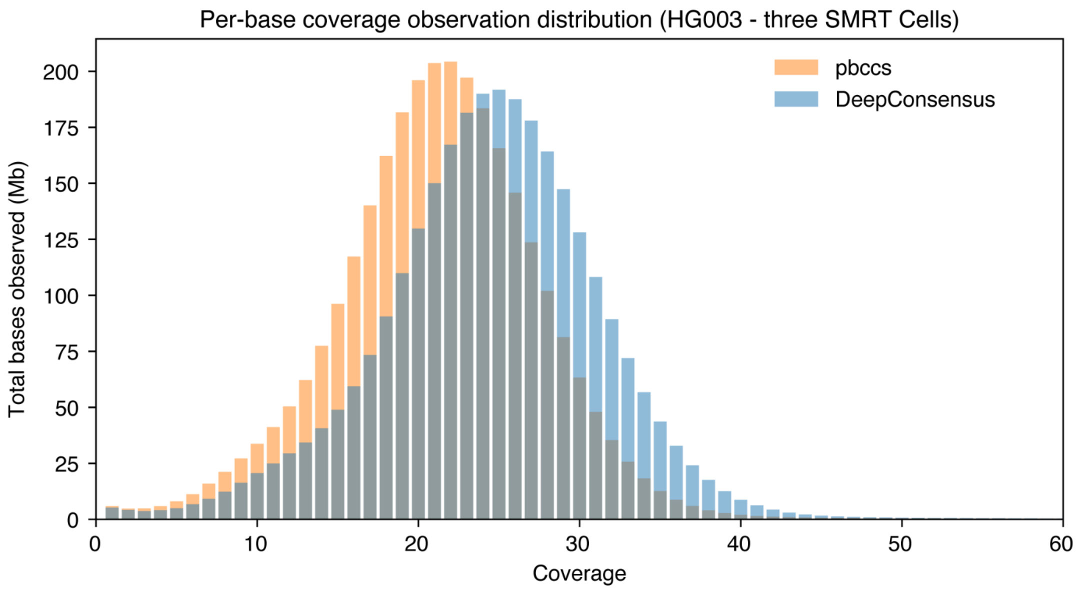


**Supplementary Figure 5** Distribution of observed coverage  of HG003 two SMRT Cells reads from pbccs and DeepConsensus.

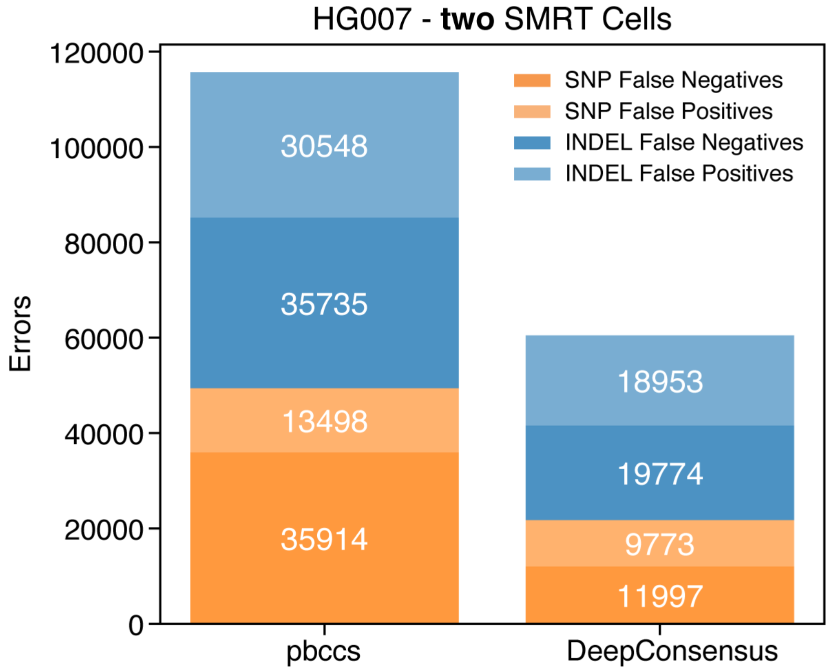


**Supplementary Figure 6** Variant calling performance with pbccs and deepconsensus reads on HG007 sample.

### Supplementary tables

**Supplementary table 1: DeepConsensus and pbccs yield for reads at Q20 (predicted 99.9% accuracy) or higher.**

| Sample | Insert size | SMRT cells | Yield at Q20 read quality | |
|---|---|---|---|---|
| | | | pbccs | DeepConsensus |
| HG003 | 15 kb | 2 | 46.15 Gb | 53.02 Gb |
| | | 3 | 65.62 Gb | 75.41 Gb |
| HG004 | 15 kb | 2 | 45.55 Gb | 52.28 Gb |
| | | 3 | 64.70 Gb | 74.26 Gb |
| HG006 | 15kb | 2 | 61.10 Gb | 69.24 Gb |
| | | 3 | 83.17 Gb | 96.56 Gb |
| HG007 | 15kb | 2 | 39.56 Gb | 49.54 Gb |

**Supplementary table 2: QUAST reported assembly statistics of hifiasm assemblies with pbccs and DeepConsensus reads.**

| Sample | SMRT cells | Method | N50 (Mb) | NG50 (Mb) | Assembly size (Gb) | Assembly completeness against GRCh38 (%) |
|---|---|---|---|---|---|---|
| HG003 15kb | 2 | pbccs | 4.74 | 4.91 | 3.16 | 97.67 |
| | | DeepConsensus | 17.23 | 17.23 | 3.1 | 97.62 |
| | 3 | pbccs | 33.64 | 33.64 | 3.1 | 97.88 |
| | | DeepConsensus | 55.54 | 55.54 | 3.1 | 97.94 |
| HG004 15kb | 2 | pbccs | 3.43 | 3.72 | 3.23 | 97.31 |
| | | DeepConsensus | 12.76 | 12.37 | 3.07 | 97.34 |
| | 3 | pbccs | 36.24 | 36.24 | 3.07 | 97.44 |
| | | DeepConsensus | 41.07 | 41.07 | 3.06 | 97.47 |
| HG006 15kb | 2 | pbccs | 18.89 | 18.55 | 3.01 | 97.37 |
| | | DeepConsensus | 31.68 | 31.54 | 3.02 | 97.5 |
| | 3 | pbccs | 42.87 | 41.19 | 3.04 | 97.76 |
| | | DeepConsensus | 51.68 | 51.68 | 3.04 | 97.88 |
| HG007 15kb | 2 | pbccs | 1.89 | 1.94 | 3.15 | 97.1 |
| | | DeepConsensus | 8.52 | 8.48 | 3.07 | 97.36 |

**Supplementary table 3: YAK estimated phred-scale Q value of hifiasm assemblies with PBCCS and DeepConsensus reads.**

| Sample | SMRT cells | Method | HAP1 Q | HAP2 Q |
|---|---|---|---|---|
| HG003 15kb | 2 | pbccs | 43.391 | 43.066 |
| | | DeepConsensus | 45.107 | 45.181 |
| | 3 | pbccs | 48.68 | 48.569 |
| | | DeepConsensus | 50.281 | 50.425 |
| HG004 15kb | 2 | pbccs | 43.446 | 43.096 |
| | | DeepConsensus | 44.916 | 44.925 |
| | 3 | pbccs | 48.881 | 48.547 |
| | | DeepConsensus | 50.333 | 50.404 |
| HG006 15kb | 2 | pbccs | 45.778 | 45.806 |
| | | DeepConsensus | 46.637 | 46.503 |
| | 3 | pbccs | 46.972 | 46.868 |
| | | DeepConsensus | 47.421 | 47.483 |
| HG007 15kb | 2 | pbccs | 39.219 | 39.067 |
| | | DeepConsensus | 42.052 | 41.935 |

**Supplementary Table 4: Assembly-based small variant calling SNP evaluation of hifiasm assemblies with pbccs and DeepConsensus reads.**

| Sample | SMRT cells | Method | SNPs | | | | | |
|---|---|---|---|---|---|---|---|---|
| | | | True positives | False negatives | False positives | Precision | Recall | F1-score |
| HG003 15kb | 2 | pbccs | 3011035 | 316460 | 31051 | 0.9049 | 0.9898 | 0.9454 |
| | | DeepConsensus | 3141265 | 186230 | 28454 | 0.9440 | 0.9910 | 0.9670 |
| | 3 | pbccs | 3235037 | 92458 | 9978 | 0.9722 | 0.9969 | 0.9844 |
| | | DeepConsensus | 3262865 | 64630 | 8834 | 0.9806 | 0.9973 | 0.9889 |
| HG004 15kb | 2 | pbccs | 2902441 | 444169 | 34944 | 0.8673 | 0.9881 | 0.9238 |
| | | DeepConsensus | 3110169 | 236441 | 31955 | 0.9293 | 0.9898 | 0.9586 |
| | 3 | pbccs | 3220904 | 125706 | 12159 | 0.9624 | 0.9962 | 0.9790 |
| | | DeepConsensus | 3268274 | 78336 | 12173 | 0.9766 | 0.9963 | 0.9863 |
| HG006 15kb | 2 | pbccs | 2898181 | 155479 | 10382 | 0.9491 | 0.9964 | 0.9722 |
| | | DeepConsensus | 2974372 | 79288 | 9682 | 0.9740 | 0.9968 | 0.9853 |
| | 3 | pbccs | 2997519 | 56141 | 8518 | 0.9816 | 0.9972 | 0.9893 |
| | | DeepConsensus | 3012599 | 41061 | 8780 | 0.9866 | 0.9971 | 0.9918 |
| HG007 15kb | 2 | pbccs | 2463514 | 605893 | 47115 | 0.8026 | 0.9813 | 0.8830 |
| | | DeepConsensus | 2739926 | 329481 | 41460 | 0.8927 | 0.9851 | 0.9366 |

**Supplementary Table 5: Assembly-based small variant calling INDEL evaluation of hifiasm assemblies with pbccs and DeepConsensus reads.**

| Sample | SMRT cells | Method | INDELs | | | | | |
|---|---|---|---|---|---|---|---|---|
| | | | True positives | False negatives | False positives | Precision | Recall | F1-score |
| HG003 15kb | 2 | pbccs | 439410 | 65091 | 221210 | 0.8710 | 0.6725 | 0.7590 |
| | | DeepConsensus | 462380 | 42121 | 128197 | 0.9165 | 0.7888 | 0.8478 |
| | 3 | pbccs | 479075 | 25426 | 48570 | 0.9496 | 0.9108 | 0.9298 |
| | | DeepConsensus | 485407 | 19094 | 25203 | 0.9622 | 0.9523 | 0.9572 |
| HG004 15kb | 2 | pbccs | 425671 | 84848 | 218034 | 0.8338 | 0.6686 | 0.7421 |
| | | DeepConsensus | 460158 | 50361 | 145983 | 0.9014 | 0.7656 | 0.8279 |
| | 3 | pbccs | 478488 | 32031 | 56204 | 0.9373 | 0.8981 | 0.9173 |
| | | DeepConsensus | 488804 | 21715 | 32312 | 0.9575 | 0.9400 | 0.9487 |
| HG006 15kb | 2 | pbccs | 365582 | 29145 | 53512 | 0.9262 | 0.8749 | 0.8998 |
| | | DeepConsensus | 375942 | 18785 | 30728 | 0.9524 | 0.9261 | 0.9391 |
| | 3 | pbccs | 377756 | 16971 | 23203 | 0.9570 | 0.9434 | 0.9502 |
| | | DeepConsensus | 381352 | 13375 | 13385 | 0.9661 | 0.9669 | 0.9665 |
| HG007 15kb | 2 | pbccs | 292676 | 104428 | 506811 | 0.7370 | 0.3706 | 0.4932 |
| | | DeepConsensus | 339622 | 57483 | 233183 | 0.8552 | 0.5982 | 0.7040 |

**Supplementary table 6: Asmgene single-copy gene completeness analysis of hifiasm assemblies with pbccs and DeepConsensus reads.**

| Sample | SMRT cells | Method | # Single copy ref | Single copy | | False duplications | | Complete (%) | Duplicated (%) |
|---|---|---|---|---|---|---|---|---|---|
| | | | | Hap1 | Hap2 | Hap1 | Hap2 | | |
| HG003 15kb | 2 | pbccs | 35374 | 32045 | 31682 | 623 | 375 | 90.076 | 1.411 |
| | | DeepConsensus | 35374 | 32882 | 32864 | 301 | 203 | 92.93 | 0.712 |
| | 3 | pbccs | 35374 | 33852 | 33668 | 180 | 141 | 95.437 | 0.454 |
| | | DeepConsensus | 35374 | 34201 | 34302 | 126 | 134 | 96.827 | 0.368 |
| HG004 15kb | 2 | pbccs | 35374 | 30973 | 30779 | 1250 | 647 | 87.284 | 2.681 |
| | | DeepConsensus | 35374 | 32306 | 31951 | 279 | 202 | 90.825 | 0.68 |
| | 3 | pbccs | 35374 | 33512 | 33183 | 225 | 142 | 94.271 | 0.519 |
| | | DeepConsensus | 35374 | 33897 | 33898 | 145 | 140 | 95.826 | 0.403 |
| HG006 15kb | 2 | pbccs | 35374 | 33149 | 33145 | 184 | 143 | 93.704 | 0.462 |
| | | DeepConsensus | 35374 | 33893 | 33765 | 186 | 143 | 95.632 | 0.465 |
| | 3 | pbccs | 35374 | 34134 | 34021 | 146 | 139 | 96.335 | 0.403 |
| | | DeepConsensus | 35374 | 34399 | 34341 | 141 | 122 | 97.162 | 0.372 |
| HG007 15kb | 2 | pbccs | 35374 | 29284 | 29056 | 649 | 453 | 82.462 | 1.558 |
| | | DeepConsensus | 35374 | 30935 | 30925 | 309 | 226 | 87.437 | 0.756 |

**Supplementary table 7: Asmgene multi-copy gene completeness analysis of hifiasm assemblies with pbccs and DeepConsensus reads.**

| Sample | SMRT cells | Method | | Multi copy | | Complete multi copy (%) | Missing multi copy (%) |
|---|---|---|---|---|---|---|---|
| | | | # Multi copy ref | Hap1 | Hap2 | | |
| HG003 15kb | 2 | pbccs | 1253 | 968 | 936 | 75.978 | 24.022 |
| | | DeepConsensus | 1253 | 980 | 982 | 78.292 | 21.708 |
| | 3 | pbccs | 1253 | 1006 | 983 | 79.37 | 20.63 |
| | | DeepConsensus | 1253 | 988 | 989 | 78.891 | 21.109 |
| HG004 15kb | 2 | pbccs | 1253 | 932 | 959 | 75.459 | 24.541 |
| | | DeepConsensus | 1253 | 967 | 942 | 76.177 | 23.823 |
| | 3 | pbccs | 1253 | 1007 | 974 | 79.05 | 20.95 |
| | | DeepConsensus | 1253 | 993 | 987 | 79.01 | 20.99 |
| HG006 15kb | 2 | pbccs | 1253 | 964 | 995 | 78.172 | 21.828 |
| | | DeepConsensus | 1253 | 1019 | 957 | 78.851 | 21.149 |
| | 3 | pbccs | 1253 | 1003 | 995 | 79.729 | 20.271 |
| | | DeepConsensus | 1253 | 1042 | 972 | 80.367 | 19.633 |
| HG007 15kb | 2 | pbccs | 1253 | 886 | 851 | 69.314 | 30.686 |
| | | DeepConsensus | 1253 | 920 | 951 | 74.661 | 25.339 |

**Supplementary Table 8: Whole genome variant calling results for DeepConsensus and pbccs reads.**

| Sample | SMRT Cells | Method | Type | Recall | Precision | F1 Score | False Negatives | False Positives |
|---|---|---|---|---|---|---|---|---|
| HG003 15kb | 2 | pbccs | INDEL | 0.969544 | 0.977546 | 0.973528 | 15365 | 11674 |
| | | | SNP | 0.994683 | 0.998822 | 0.996748 | 17693 | 3907 |
| | | DeepConsensus | INDEL | 0.976595 | 0.978256 | 0.977425 | 11808 | 11383 |
| | | | SNP | 0.996891 | 0.99904 | 0.997964 | 10345 | 3191 |
| | | DeepConsensus + DeepVariant v1.2 | INDEL | 0.941865 | 0.937756 | 0.939806 | 29329 | 32591 |
| | | | SNP | 0.996948 | 0.999033 | 0.997989 | 10155 | 3214 |
| HG003 15kb | 3 | pbccs | INDEL | 0.986755 | 0.988421 | 0.987587 | 6682 | 6069 |
| | | | SNP | 0.998211 | 0.999261 | 0.998736 | 5954 | 2457 |
| | | DeepConsensus | INDEL | 0.988248 | 0.988042 | 0.988145 | 5929 | 6278 |
| | | | SNP | 0.998487 | 0.999382 | 0.998934 | 5035 | 2056 |
| | | DeepConsensus + DeepVariant v1.2 | INDEL | 0.956454 | 0.954828 | 0.95564 | 21969 | 23606 |
| | | | SNP | 0.99856 | 0.999318 | 0.998939 | 4791 | 2268 |
| HG004 15kb | 2 | pbccs | INDEL | 0.966677 | 0.976135 | 0.971383 | 17012 | 12540 |
| | | | SNP | 0.992949 | 0.99839 | 0.995662 | 23596 | 5363 |
| | | DeepConsensus | INDEL | 0.9746 | 0.977708 | 0.976152 | 12967 | 11797 |
| | | | SNP | 0.995903 | 0.998765 | 0.997332 | 13711 | 4123 |
| HG004 15kb | 3 | pbccs | INDEL | 0.98602 | 0.988074 | 0.987046 | 7137 | 6327 |
| | | | SNP | 0.99782 | 0.999242 | 0.99853 | 7297 | 2535 |
| | | DeepConsensus | INDEL | 0.988143 | 0.988352 | 0.988248 | 6053 | 6190 |
| | | | SNP | 0.998234 | 0.999352 | 0.998793 | 5910 | 2168 |
| HG006 15kb | 2 | pbccs | INDEL | 0.984686 | 0.985165 | 0.984925 | 6215 | 6207 |
| | | | SNP | 0.997865 | 0.996777 | 0.997321 | 6366 | 9632 |
| | | DeepConsensus | INDEL | 0.989101 | 0.987795 | 0.988448 | 4423 | 5115 |
| | | | SNP | 0.998551 | 0.997181 | 0.997865 | 4322 | 8426 |
| HG006 15kb | 3 | pbccs | INDEL | 0.989377 | 0.986936 | 0.988155 | 4311 | 5484 |
| | | | SNP | 0.99896 | 0.996987 | 0.997973 | 3101 | 9012 |
| | | DeepConsensus | INDEL | 0.993978 | 0.991333 | 0.992653 | 2444 | 3639 |
| | | | SNP | 0.999297 | 0.997277 | 0.998286 | 2098 | 8145 |

**Supplementary table 9. Yield for 15kb and 24kb reads at Q20 used for assembly and variant calling experiments.**

| Sample | Insert size | SMRT cells | Method | Yield |
|---|---|---|---|---|
| HG002 | 15 kb | 2 | DeepConsensus | 88.98 Gb |
| | 24 kb | 2 | DeepConsensus | 103.61 Gb |

**Supplementary table 10: QUAST reported assembly statistics of hifiasm assemblies with 15kb and 24kb reads.**

| Sample | Read length | SMRT cells | Method | N50 (Mb) | NG50 (Mb) | Assembly size (Gb) | Assembly completeness against GRCh38 |
|---|---|---|---|---|---|---|---|
| HG002 | 15kb | 2 | DeepConsensus | 24.81 | 24.81 | 3.09 | 97.654 |
| | 24kb | 2 | DeepConsensus | 33.14 | 34.05 | 3.13 | 97.68 |

**Supplementary table 11: YAK estimated phred-scale Q value of hifiasm assemblies with 15kb and 24kb reads.**

| Sample | Read length | SMRT cells | Method | HAP1 Q | HAP2 Q |
|---|---|---|---|---|---|
| HG002 | 15kb | 2 | DeepConsensus | 51.879 | 51.534 |
| | 24kb | 2 | DeepConsensus | 50.901 | 50.614 |

**Supplementary Table 12: Assembly-based small variant calling SNP evaluation of hifiasm assemblies with 15kb and 24kb reads.**

| Sample | Read length | SMRT cells | Method | SNPs | | | | | |
|---|---|---|---|---|---|---|---|---|---|
| | | | | True positives | False negatives | False positives | Precision | Recall | F1-score |
| HG002 | 15kb | 2 | DeepConsensus | 3318960 | 46167 | 7073 | 0.986281 | 0.997876 | 0.992045 |
| | 24kb | 2 | DeepConsensus | 3312496 | 52631 | 7404 | 0.9844 | 0.9978 | 0.9910 |

**Supplementary Table 13: Assembly-based small variant calling INDEL evaluation of hifiasm assemblies with 15kb and 24kb reads.**

| Sample | Read length | SMRT cells | Method | INDELs | | | | | |
|---|---|---|---|---|---|---|---|---|---|
| | | | | True positives | False negatives | False positives | Precision | Recall | F1-score |
| HG002 | 15kb | 2 | DeepConsensus | 507578 | 17891 | 19463 | 0.965952 | 0.964251 | 0.965101 |
| | 24kb | 2 | DeepConsensus | 511634 | 13835 | 19131 | 0.9737 | 0.9652 | 0.9694 |

**Supplementary table 14: Asmgene single-copy gene completeness analysis of hifiasm assemblies with 15kb and 24kb reads.**

| Sample | Read length | SMRT cells | Method | # Single copy ref | Single copy | | False duplications | | Complete (%) | Duplicated (%) |
|---|---|---|---|---|---|---|---|---|---|---|
| | | | | | Hap1 | Hap2 | Hap1 | Hap2 | | |
| HG002 | 15kb | 2 | DeepConsensus | 35374 | 34553 | 34353 | 153 | 114 | 97.396 | 0.377 |
| | 24kb | 2 | DeepConsensus | 35374 | 34517 | 34269 | 191 | 189 | 97.227 | 0.537 |

**Supplementary table 15: Asmgene multi-copy gene completeness analysis of hifiasm assemblies with 15kb and 24kb reads.**

| Sample | Read length | SMRT cells | Method | # Multi copy ref | Multi copy | | Complete multi copy (%) | Missing multi copy (%) |
|---|---|---|---|---|---|---|---|---|
| | | | | | Hap1 | Hap2 | | |
| HG002 | 15kb | 2 | DeepConsensus | 1253 | 1024 | 904 | 76.935 | 23.065 |
| | 24kb | 2 | DeepConsensus | 1253 | 1043 | 985 | 80.926 | 19.074 |

**Supplementary Table 16: Chr20 variant calling results for DeepConsensus reads.**

| Sample | SMRT Cells | Method | Type | Recall | Precision | F1 Score | False Negatives | False Positives |
|---|---|---|---|---|---|---|---|---|
| HG002 15kb | 2 | DeepConsensus | INDEL | 0.989606 | 0.987434 | 0.988518 | 117 | 147 |
| | | | SNP | 0.998822 | 0.999748 | 0.999285 | 84 | 18 |
| HG002 24kb | 2 | DeepConsensus | INDEL | 0.989961 | 0.989044 | 0.989502 | 113 | 128 |
| | | | SNP | 0.999257 | 0.999734 | 0.999495 | 53 | 19 |
